# TMVisDB: Annotation and 3D-visualization of transmembrane proteins

**DOI:** 10.1101/2024.11.22.624323

**Authors:** Tobias Olenyi, Céline Marquet, Anastasia Grekova, Leen Houri, Michael Heinzinger, Christian Dallago, Burkhard Rost

## Abstract

Since the rise of cellular life, transmembrane proteins (TMPs) have been crucial to various cellular processes through their central role as gates and gatekeepers. Despite their importance, experimental high-resolution structures for TMPs remain underrepresented due to experimental challenges. Given its performance leap, structure predictions have begun to close the gap. However, identifying the membrane regions and topology in three-dimensional structure files on a large scale still requires additional *in silico* predictions. Here, we introduce *TMVisDB* to sieve through millions of predicted structures for TMPs. This resource enables both browsing through 46 million predicted TMPs and visualizing them along with their topological annotations without having to tap into costly predictions of the AlphaFold3-style. *TMVisDB* joins *AlphaFoldDB* structure predictions and transmembrane topology predictions from the protein language model (pLM) based method *TMbed*. We showcase the utility of TMVisDB for the analysis of proteins through two use cases, namely the B-lymphocyte antigen CD_20_ (*Homo sapiens*) and the cellulose synthase (*Novosphingobium sp. P6W*). We demonstrate the value of *TMVisDB* for large-scale analyses through findings pertaining to all TMPs predicted for the human proteome. TMVisDB is freely available at https://tmvisdb.rostlab.org

## Introduction

### Experimental structures for transmembrane proteins remain sparse

Despite many biological processes depending on proteins spanning cell membranes, only 5% of all high-resolution experimental three-dimensional (3D) structures in the Protein Data Bank (PDB) are transmembrane proteins (TMPs) [6]. Although structure-determination methods have advanced significantly [1, 7-11], TMPs continue to pose many technical challenges [12]. Therefore, today’s databases often contain only small subsets of membrane proteins, and few focus solely on TMPs or add information on (predicted) membrane embeddings [13-17]. As of November 2024, *mpstruc*, a curated database, covers 1,736 unique TMPs with experimental 3D structures [18], constituting 1.2% of all PDB structures (132,007 unique sequences, defined as: no pair has 100% percentage pairwise sequence identity). Substantially below the estimate of 20-30% of proteins in any proteome being TMPs [2, 19], this imbalance becomes more striking, considering that TMPs make up almost 50% of all drug targets [20].

### New resources may push novelty into the realm of TMPs

Few computational methods have revolutionized a field as quickly and dramatically as AlphaFold2 [1]. Providing reliable 3D predictions, AlphaFold2, AlphaFoldDB (AFDB) [3], and AlphaFold3 [7] are extraordinary catalysts for research in artificial intelligence (AI) and the life sciences [21]. Along with the advance in structure prediction, methods of quickly and reliably searching and clustering the exploding sequences have continuously improved [22-24]. An orthogonal approach to leverage the wealth of sequence data is provided by protein Language Models (pLMs) [25-30]. As a special case of Large Language Models (LLMs) as have been popularized by ChatGPT, pLMs learn from protein sequences without experimental annotations (dubbed ‘unlabeled’ in AI jargon) to extract meaningful numeric representations (embeddings) encoding sequences as vectors. The knowledge acquired during pre-training on raw sequences can readily be transferred to downstream predictions of per-residue [2, 31-34] (e.g., which residue is in a membrane helix) and per-protein phenotypes [35-38] (e.g. TMP or not).

The unparalleled number of available protein sequences, pLM embeddings, and experimental and predicted structures may help to overcome limitations for studying TMPs *in vitro* and *in vivo*. Combining *in silico* methods and resources enables the discovery of previously unknown TMPs and the exploration of known and unknown TMPs in exceptional detail. An example of the usefulness of TMP predictions is the *TmAlphaFold database*, which takes AlphaFold2 structures as input to provide membrane localization for almost 216k (215,844) predicted alpha-helical TMPs [39].

Here, we introduce TMVisDB, a collection of over 46 million (46M) predicted alpha-helical and beta-barrel TMPs with their AlphaFold2-predicted 3D structure as well as membrane topology and annotation predicted by the pLM-based method TMbed [2]. TMVisDB was created by applying TMbed to all monomeric proteins in AFDB [3] (>200M, 07/22). Most of these proteins are unreviewed, i.e., their sequences are not experimentally verified [5]. TMbed is *on par* with or outperforms all other TMP prediction methods in classifying residues as transmembrane alpha-helix (TMH) or transmembrane beta-strand (TMB). This state-of-the-art (SOTA) corresponds to correctly predicting all transmembrane segments (TMH or TMB) for about 90% of all proteins. The measure for “correct prediction” is an overlap of ±5 residues between the predicted and observed segments. Although this is slightly less accurate than the difference between different assignments [40-42], mostly the errors are on one side only, i.e., the prediction is shifted because most TM segments are predicted similar in length as the ones observed. TMbed also predicts the orientation of TM segments within the membrane (dubbed *topology*) and signal peptides (SP). The latter reaches the binary classification performance of SignalP6 without distinguishing different SP types [2, 43]. TMVisDB is the first resource to gather such a large number of per-residue transmembrane topology annotations and, compared to existing TMP databases [18, 39], expanding the exploration space of easily browsable TMPs by two to four orders of magnitude. Providing interactive visualizations of TMP annotations with their respective AlphaFold2 structures, the intuitive interface of tmvisdb.rostlab.org enables all users to investigate small- and large-scale hypotheses on TMPs.

## Results

### Over 46M predicted transmembrane proteins

TMVisDB is a relational database with 46,048,450 proteins (11/24, Table 1), comprising 23% of AlphaFold DB [3] (AFDB, >200M, 07/22) [21]. Transmembrane proteins (TMPs) and their topology were predicted for proteins in AFDB using the pLM-based, state-of-the-art TMP prediction method TMbed [2] and discarding proteins without a predicted transmembrane section. About 95% of the proteins in TMVisDB have predicted transmembrane alpha-helices (TMHs), 4.6% contain predicted transmembrane beta-strands (TMBs), and about 1‰ (0.1%) are predicted to contain both, likely providing a lower estimate for the error of our approach as these predictions are likely wrong (although this continues to be an assumption). TMbed predicted signal peptides (SP) for 13.6% of all TMPs (archaea: 13.83%, bacteria: 14.5%, eukaryotes: 15.3%). Taking all proteins with AFDB entries (TMP and non-TMP), TMbed predicted about 7% to have signal peptides. Despite differences in the underlying datasets, this prediction rate aligns with SignalP6’s published estimate of 8% for eukaryotic proteins [43].

**Table 1:**
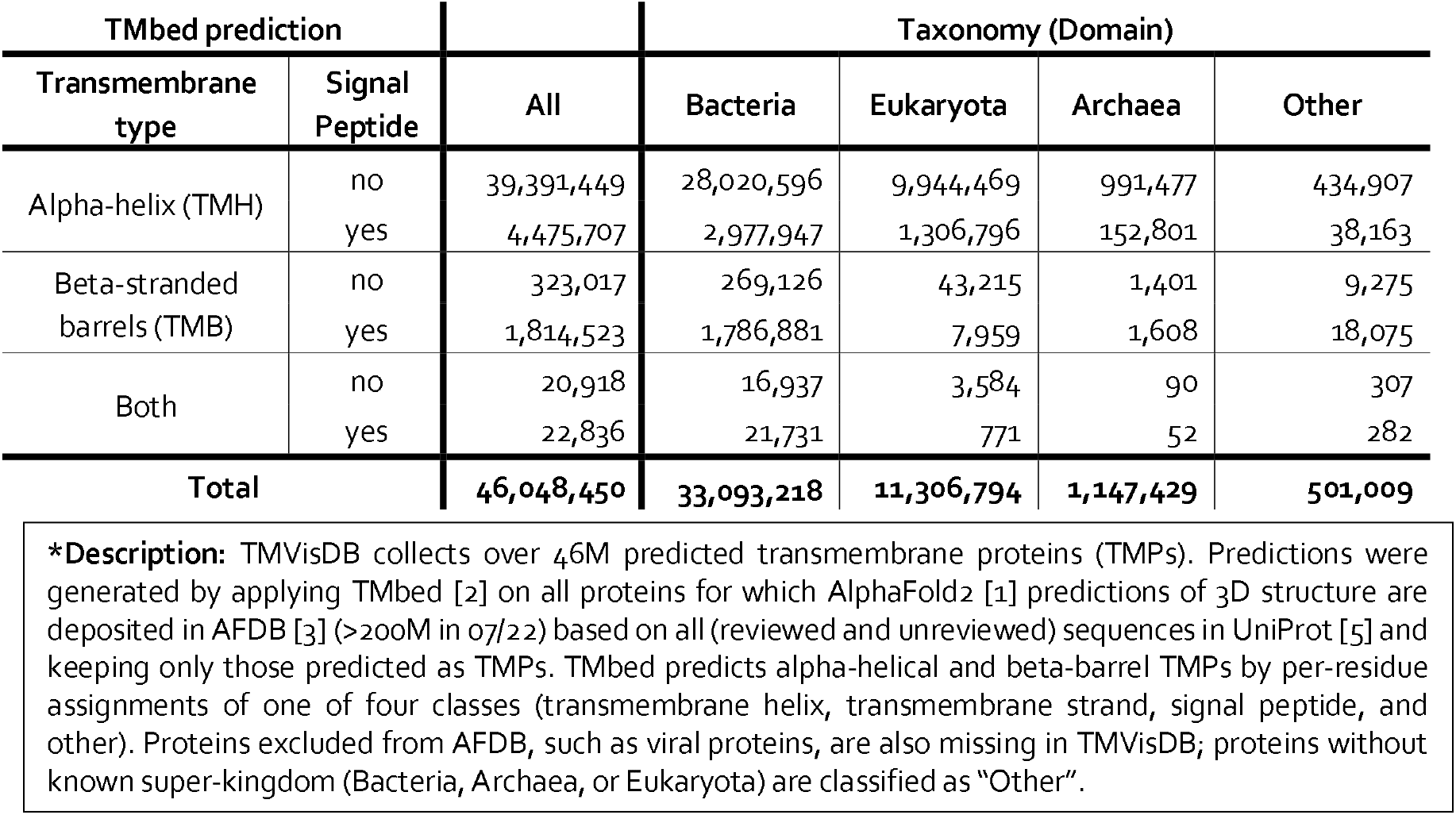
Summary of proteins in TMVisDB*.

### Intuitive browsing and visualization

The website tmvisdb.rostlab.org enables interactive access to browse and visualize all TMPs in TMVisDB. For large-scale analyses, the database can be filtered by taxonomy (UniProt organism identifier, domain, kingdom), predicted TMP-type (TMH, or TMB, or both), protein length, and presence/absence of predicted SP (Fig. 1a). Each database record contains: UniProt name (ID), sequence length, organism domain and kingdom, a link to the organism on UniProtKB, sequence classification based on TMbed predictions, and cumulative TMbed predictions [2]. Results are downloadable in CSV format (comma-separated columns in ASCII). Each row in the results is clickable to visualize the prediction details, corresponding AlphaFold2 3D structure, and available annotations. In the detail view, the structure of each protein can be colored by the (1) available annotations or (2) AlphaFold2 per-residue confidence metric (pLDDT; Fig. 2, Supporting Online Material (SOM) Fig. S1 for the interface, S2-S4). Protein sequence annotations are shown below the 3D visualization, along with links for the respective sources; possible annotation resources are UniProtKB [5], Membranome [13], TMAlphaFold [39], TopDB [44], and TMbed predictions [2]. Individual annotation regions are clickable and trigger highlighting of the respective region in the 3D visualization.

**Fig. 1:**
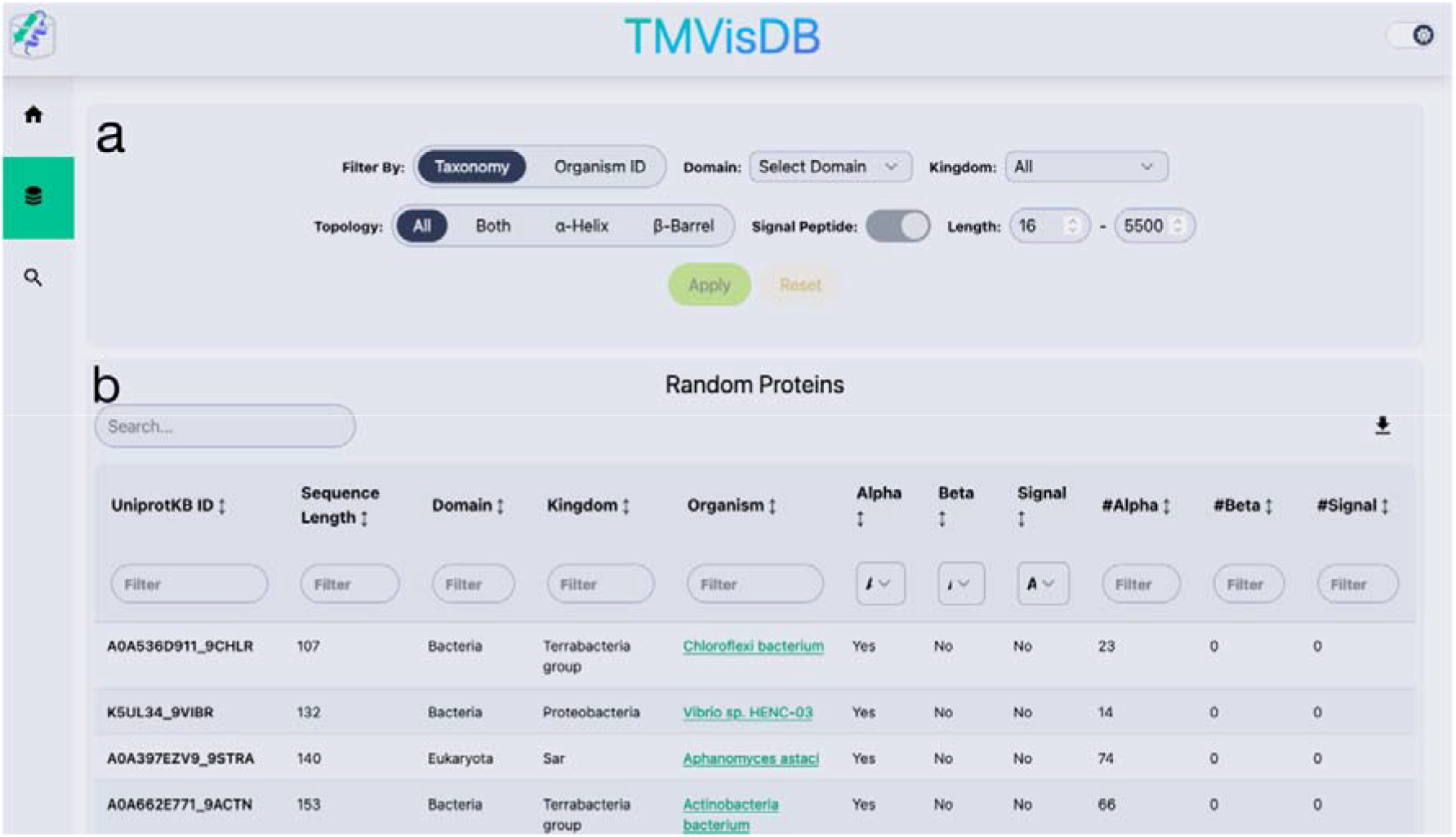
TMVisDB filter interface. The interface has two main components. Panel **(a)** displays the primary filtering system with four distinct categories: (1) Taxonomy, allowing selection by UniProt Organism Identifier, Domain, and Kingdom; (2) Transmembrane topology, distinguishing between alpha-helix and beta-strand structures; (3) Signal Peptides (SP), enabling inclusion or exclusion of sequences with predicted SP; and (4) Sequence length parameters. Panel **(b)**: presents a partial output from a random TMVisDB selection without applied filters. Each record contains seven key features: (1) UniProtKB name (ID), (2) sequence length, (3) organism domain, (4) kingdom classification, (5) organism link to UniProtKB, (6) sequence classification based on TMbed predictions[2], and (7) cumulative TMbed predictions [2, 4].

**Fig. 2:**
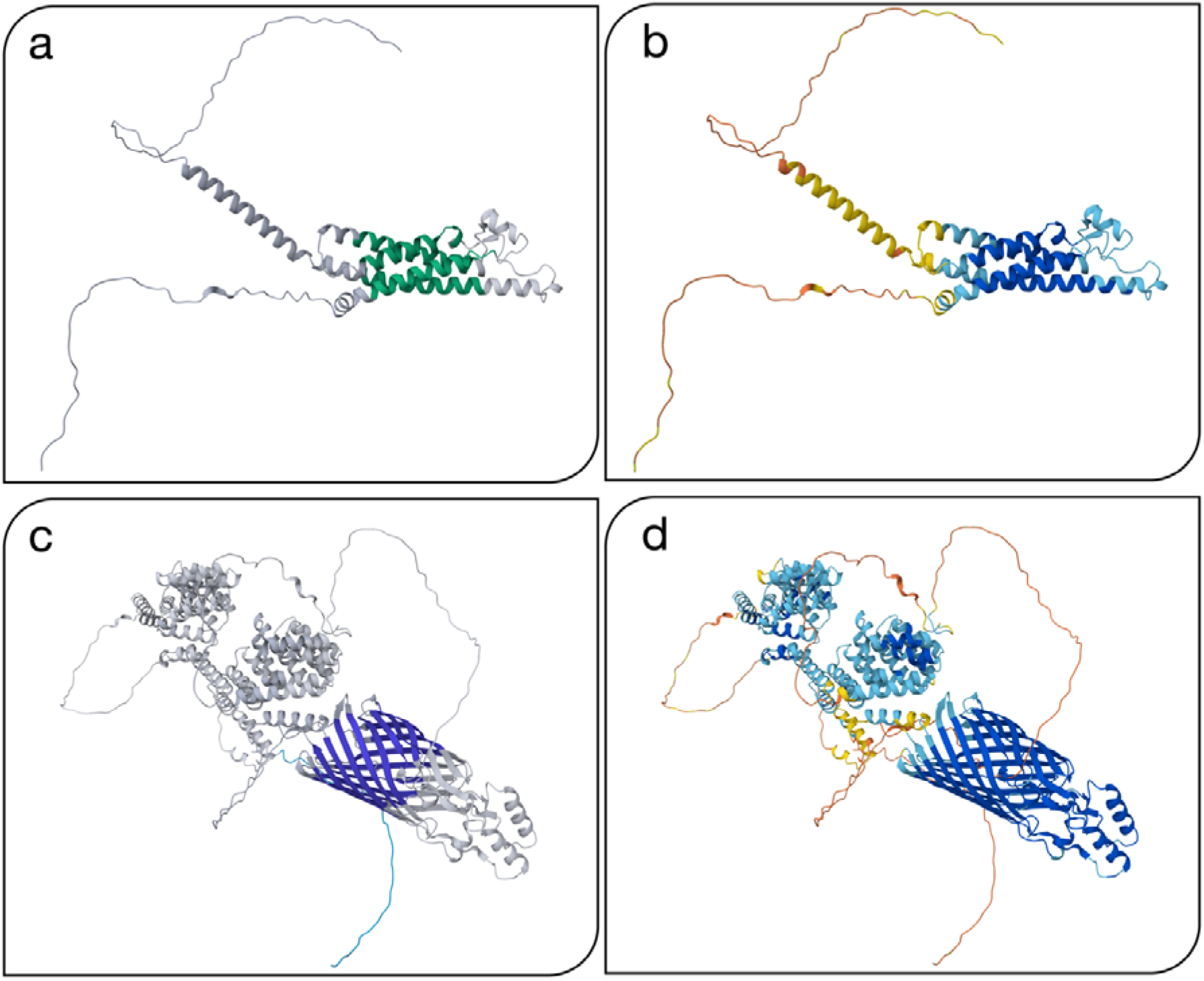
TMVisDB single protein use cases. Structural analysis of two proteins: (1) *B-lymphocyte antigen CD20* (P11836, Homo sapiens) and (2) *cellulose synthase protein* (A0A2Z5E6S1, Novosphingobium sp. P6W) using AlphaFold2 prediction [1]. Panels show CD20 in the upper section **(a**, **b)** and cellulose synthase protein in the lower section (c, d). Left panels **(a**, **c)** visualize TMbed’s predicted transmembrane topology using three distinct colors: (1) blue for alpha-helix, (2) gray for non-membrane regions, and (3) turquoise for signal peptide. Right panels **(b**, **d)** display AlphaFold2’s per-residue confidence metric (pLDDT) with four confidence levels: (1) very confident (pLDDT > 90, blue), (2) confident (70 < pLDDT ≤ 90, green), (3) low confidence (50 < pLDDT ≤ 70, yellow), and (4) very low confidence (pLDDT ≤ 50, red). The predicted transmembrane regions exhibit strong colocalization with high structural confidence, as indicated by high pLDDT scores. Non-membrane regions, particularly those with loop conformations, display lower confidence scores, consistent with their inherent flexibility.

### Use case 1: B-lymphocyte antigen CD20 - expanding annotations

The B-lymphocyte antigen CD20 (UniProtKB-ID: P11836, Fig. 2a,b) is an ideal use case. Although, well-studied nature and an important drug target, there still is no high-resolution experimental structure available for CD20 in isolation.

TMVisDB revealed four transmembrane helices (TMHs, with 18-24 residues), supporting the high-confidence AlphaFold2 predictions (Fig. 2b). The predicted topology largely confirms the membrane orientation observed in existing experimental structures [45] with minor deviations of up to five residues between TMVisDB and UniProtKB [46].

Our predictions also reinforce CD_20_’s classification as a member of the MS4A family of TMPs. The structural features align with previous studies [47, 48], showing a non-glycosylated tetraspanin structure with one shorter and one longer extracellular loop and both C- and N-termini located in the cytosol (Fig. 2a). This case study demonstrates how TMVisDB can provide valuable structural insights for important therapeutic targets, even in the absence of complete experimental structures.

### Use case 2: Cellulose synthase - exploring novel TMP annotation

TMVisDB’s capability to analyze previously unstudied transmembrane proteins is exemplified by our examination of the *cellulose synthase protein* (UniProtKB-ID: A0A2Z5E6S1) from *Novosphingobium sp. P6W*, a gram-negative bacterium (Fig. 2c, d). Based on UniProtKB’s automatic annotation of unreviewed records, this protein localizes to the prokaryotic outer membrane and functions in glycan metabolism and bacterial cellulose biosynthesis [49].

Our analysis revealed significant agreement between multiple prediction methods. While UniProtKB annotated a signal peptide spanning 29 residues, TMbed extended this prediction to 40 residues, confirmed by a high-confidence prediction from SignalP6.0 (cleavage site at position 40, probability 0.97) [43]. TMbed additionally identified twelve transmembrane beta-strands (TMBs), which correspond closely to the beta-barrel configuration predicted by AlphaFold2 (Fig. 2c). The high pLDDT scores of the transmembrane residues provide further validation for these previously unknown TMBs (Fig. 2d).

We observed some minor discrepancies in the predictions, particularly where certain TMBs appeared shorter than their corresponding segments in the AlphaFold2 barrel structure (Fig. 2c, lower right). Several hypotheses could explain these differences: the natural membrane embedding might influence segment length, TMbed may have produced false negative predictions for some residues, or protein dynamics during binding events could have led AlphaFold2 to predict an alternative conformational state. However, current evidence remains insufficient to support any of these explanations conclusively.

### Use case 3: *Homo sapiens* - combining large- and small-scale analyses

In addition to studies of individual proteins, TMVisDB enables large-scale analyses. TMVisDB contains 20.7% of the human proteins in AFDB (11/24), accessible via the human UniProtKB organism identifier (no. 9606). Out of these, we investigated the only proteins for which TMbed predicted both TMH and TMB: (1) *Deoxyribonuclease I fragment* (UniProtKB-ID: A0A192ZHB2; Fig. S2), (2) two sequence-similar *perforins* (UniProtKB-ID: B3KUT2, Q2M385; Fig. S3), (3) and three sequence-similar *DnaJ* homologs (B4DGD5, B4DPK2, Q9NVH1; Fig. S4).

TMbed predicted contradicting inside-to-outside TMH and outside-to-inside TMB topologies for the Deoxyribonuclease I fragment. While AlphaFold2 confirmed secondary structure elements in these regions, only the beta-strand was predicted at high plDDT (>80), and the prediction lacked an expected loop/bend between elements (Fig. S2). These inconsistencies suggest partial correctness of either prediction at best.

The perforin predictions showed mixed evidence for transmembrane topology. While the TMH prediction aligned with AlphaFold2 structure prediction (Fig. S3), UniProt annotation [50], and known membrane interactions [51], the short predicted TMB (5-6 residues) lacked such support. Although *MPEG1/Perforin-2* (UniProtKB-ID: Q2M385) forms homo-oligomeric beta-barrel pores [52], comparative analysis of membrane-targeting perforin-like proteins from *Toxoplasma gondii* (UniProtKB-ID: G3G7T0) and *Plasmodium falciparum* (UniProtKB-ID: Q9U0J9) showed no similar TMB patterns, suggesting this prediction to be incorrect.

For *DnaJ* homologs, TMbed and AlphaFold2 predictions showed remarkable agreement in TMH and TMB membrane boundary predictions (Fig. S4). These predictions align with *DnaJ*’s known role in the mitochondrial intermembrane space bridging (MIB) complex and its association with the mitochondrial contact site and cristae junction organizing system (*MICOS*) [53, 54]. While direct experimental validation of these transmembrane domains remains pending, their presence could explain *DnaJ*’s function in *MICOS* assembly and cristae organization, presenting either a novel finding or a hypothesis for experimental verification.

## Discussion

### Organizing the explosion of structures

The exponential growth of protein sequences and structures necessitates efficient organization and accessibility of specific protein groups. Protein language models (pLMs) have revolutionized this organization by enabling rapid functional scanning of large sequence databases independent of traditional sequence alignment methods [55, 56]. This capability enabled us to efficiently identify and extract transmembrane proteins (TMPs) from UniProt using the pLM-based TMbed method. While several structure prediction methods are now available [8-11], we chose AlphaFold2 [1] for TMVisDB due to its demonstrated accuracy in TMP structure prediction [57] and successful application in resources such as the TmAlphaFold database [39]. The resulting TMVisDB combines these predictions with an intuitive search and visualization interface, making TMP structural data readily accessible to researchers.

### Benefits and limitations often go hand in hand

TMVisDB enhances TMP analysis by combining per-residue transmembrane predictions with higher-resolution 3D structure predictions through complementary approaches: the pLM-based TMbed provides low-resolution protein-specific predictions, while AlphaFold2 contributes high-resolution family-averaged predictions from multiple sequence alignments (MSAs). This dual approach provides independent validation, as demonstrated by the cellulose synthase protein case, for which aligned predictions revealed uncertainty about membrane embedding depth. The B-lymphocyte antigen CD20 case further illustrates TMVisDB’s utility in confirming TM regions and its current limitations to monomers, a challenge that future developments like AlphaFold Multimer [58] might address.

TMbed, underlying TMVisDB, shines with a very low false positive rate (<1%). However, given the millions of predictions (> 46 million), this still implies many errors in our resource, as indicated by the likely incorrect prediction of proteins with both TMH and TMB segments (0.1% of entries) - a biologically improbable combination. The database enables rapid comparative analyses to identify such potential false positives, as demonstrated in our perforin analysis. However, TMVisDB faces significant scale-related challenges: it currently contains 38K human TMPs, far exceeding the estimated 20K human proteins [59]. This inflation likely stems from AFDB including unreviewed proteins and fragments, creating redundancy that, while potentially useful for some analyses, complicates database navigation.

Beyond classification accuracy, TMbed predicts transmembrane segment boundaries within ±5 residues of observed positions, often resulting in shifted rather than extended predictions. This level of positional variance is comparable to the disagreement between experimental methods [2, 40, 60], potentially reflecting methodological uncertainties in terminus assignment or differences in protein folding states [60].

Future developments of TMVisDB will improve by implementing GO annotation- and binding filters [32, 38], improving database performance, and introducing AFDB syncing. These enhancements, guided by user feedback, will strengthen TMVisDB’s role as a comprehensive resource for TMP research.

## Materials and Methods

### TMbed transmembrane topology prediction

TMbed [2] performs residue-level classification of transmembrane proteins (TMPs) into four categories: transmembrane alpha-helix (TMH), transmembrane beta-strand (TMB), signal peptide (SP), and others. The method predicts membrane orientation for TMH and TMB segments without the capability to identify re-entrant regions. TMbed achieves state-of-the-art performance with 98±1% accuracy for alpha-helical and 94±8% for beta-barrel TMPs while maintaining a false positive rate below 1%. Its predictions locate 90% of TMHs and TMBs within five residues of their actual position. TMbed can process the entire Swiss-Prot [5] (∼567,000 proteins) in under 9 hours using only protein Language Model (pLM) embeddings from single sequences, making it ideal for high-throughput applications like TMVisDB [2].

### Database and backend

TMVisDB is implemented as an SQLite (version 3) database with a FastAPI-based Python backend. Each entry is uniquely identified by its UniProt accession (UA) and contains protein sequence data, predictions, annotations, and taxonomic information. The database incorporates, where available, experimentally derived topology data from TOPDB [44] and single alpha-helix transmembrane annotations from Membranome [13]. TMVisDB includes all transmembrane proteins from AlphaFold DB (AFDB) [21], as predicted by TMbed [2]. AFDB’s sequence coverage limitations apply to TMVisDB: sequences must be (i) between 16 and 2,700 amino acids for UniProtKB/Swiss-Prot entries (1,280 for others), (ii) contain only standard amino acids, (iii) appear in UniProtKB’s reference proteome “one sequence per gene” FASTA file, (iv) remain unmodified in recent UniProt releases, and (v) be non-viral. The database and its API are hosted at the Technical University of Munich (TUM); the code is available at https://github.com/t03i/TMVisDB.

### Web frontend

The TMVisDB web interface is implemented in Svelte 4, providing an interactive environment for exploring transmembrane protein data. Users can filter the database by transmembrane topology (TMH, TMB), signal peptide presence, taxonomy, and protein length (Fig. 1). For individual proteins, the interface offers detailed visualization using the PDBe Molstar component for 3D structure display and the Nightingale feature viewer from EBI for sequence annotations (Fig. S2). The visualization system enables dynamic structures coloring based on either transmembrane topology predictions or AlphaFold2’s pLDDT scores (Fig. 2). Additional annotations from UniProtKB [5] and TMAlphaFold [39] are loaded on demand. At the same time, structural data is fetched directly from AlphaFoldDB. The web interface source code is available at https://github.com/t03i/TMVisDB; the web interface is available at tmvisdb.rostlab.org.

### Accession Numbers

UniProt accessions: <u>A0A2Z5E6S1</u>, <u>P11836</u>, <u>A0A192ZHB2</u>, <u>B3KUT2</u>, <u>Q2M385</u>, <u>B4DGD5</u>, <u>B4DPK2</u>, <u>Q9NVH1</u>, <u>G3G7T0</u>, <u>Q9U0J9</u> UniProt organism identifier: 9606

## Abbreviations

AFDB: AlphaFold DB
AI: artificial intelligence
ASCII: American Standard Code for Information Interchange
GO: Gene Ontology
MIB: mitochondrial intermembrane space bridging complex
MICOS: mitochondrial contact site and cristae junction organizing system
PDB: Protein Data Bank
pLDDT: predicted local distance difference test
pLM: protein language model
SAM: mitochondrial outer membrane sorting assembly machinery
SOM: supporting online material
SP: signal peptide
TM: transmembrane
TMB: transmembrane beta strand
TMH: transmembrane alpha helix
TMP: transmembrane protein
TOPDB: Topology Database of Transmembrane Proteins
UA: UniProt accession number
3D: three dimensional

## Data Availability

The database and web interface presented in this study are freely accessible at tmvisdb.rostlab.org. A static version of the database (excluding sequence information and indices due to size constraints) has been deposited in the Zenodo repository (<u>10.5281/zenodo.14186950</u>). The complete source code for database implementation and web interface is available through GitHub (https://github.com/t03i/TMVisDB) under the Apache 2.0 license. TMVisDB integrates data from the following established resources: AlphaFold DB (https://alphafold.ebi.ac.uk/), UniProtKB (https://www.uniprot.org/), TOPDB (http://topdb.enzim.hu/), Membranome (https://membranome.org/), and TMAlphaFold (http://tmalphafold.ttk.hu/). All data and source code are available without restrictions for academic and non-commercial use.

## Acknowledgments

We gratefully acknowledge Dr. Luisa F. Jimenez-Soto for her critical analysis of our results and Dr. Michael Bernhofer for his invaluable guidance on the project direction and manuscript preparation. We extend our appreciation to Nikita Kugut for his continued support throughout this work, and to Prof. Arne Skerra (Technical University of Munich, WZW) for suggesting the CD20 system as a valuable use case.

We thank the Technical University of Munich for providing the infrastructure to host TMVisDB. Our work has been made possible by the groundbreaking development of AlphaFold2 by Dr. John Jumper and the DeepMind team, and we appreciate their commitment to open science through the public release of both code and predictions.

This work builds upon the contributions of numerous individuals and institutions committed to open science. We acknowledge the developers of the open-source programming libraries utilized in this study, the researchers who deposit their experimental data in public databases, the dedicated teams maintaining these databases, and the scientists who develop and share methods for enriching experimental data.

## Funding

The Bavarian Ministry of Education supported this work by funding TUM. The authors declare no conflicts of interest.

## Author contributions

**Tobias Olenyi:** Conceptualization, Data Curation, Methodology, Project Administration, Software, Validation, Visualization, Writing – Original Draft, Writing – Review & Editing

**Céline Marquet:** Conceptualization, Data Curation, Project Administration, Validation, Writing – Original Draft, Writing – Review & Editing

**Anastasia Grekova:** Formal Analysis, Software, Writing – Original Draft

**Leen Houri:** Formal Analysis, Writing – Original Draft

**Michael Heinzinger:** Conceptualization, Writing – Original Draft

**Christian Dallago:** Conceptualization, Validation

**Burkhard Rost:** Supervision, Writing – Review & Editing

## Notes

https://doi.org/10.5281/zenodo.14186950

https://github.com/t03i/TMVisDB

https://tmvisdb.rostlab.org/

### Competing Interest Statement

The authors have declared no competing interest.

### Summary of Updates

Highlighted the unreviewed nature of the database content; Clarified focus on monomeric structures due to AlphaFoldDB limitations; Explained accuracy metrics for transmembrane helix predictions (+-5 residues);Clarified unique sequence definition (100% identity threshold); author affiliations updated

