## Supplementary Material for "TMVisDB: Annotation and 3D-visualization of transmembrane proteins"

### TMVisDB: Annotation and 3D-visualization of transmembrane proteins - Supporting Material

##### Short Description

This supplementary document provides additional visualizations and analyzes to complement the main manuscript. We present a screenshot of the protein detail page interface, demonstrating its features and functionality (**Fig. S1**). Additionally, we provide figures illustrating the AlphaFold2-predicted 3D structures [1] and TMbed-predicted membrane topologies [2] for the following proteins from our Homo sapiens case study:

- Deoxyribonuclease (DNase) I fragment (DNASE1) (A0A192ZHB2) (**Fig. S2**)
- Macrophage-expressed gene 1 (MPEG1/Perforin-2) (Q2M385)(**Fig. S3**)
- DnaJ homolog subfamily C member 11 (DNAJC11/HSP40) (Q2M385)(**Fig. S4**)

##### Color Scheme

Fig. S1-S3: All panels show the 3D protein structures predicted by AlphaFold2 [12]. Left panels (**a**) visualize TMbed's predicted transmembrane topology using four distinct colors: (1) green for transmembrane helices (TMH), (2) blue for transmembrane beta-barrels (TMB), (3) turquoise for signal peptides, and (4) gray for other regions. Right panels (**b**) display AlphaFold2's per-residue confidence metric (pLDDT) with four confidence levels: (1) very confident (pLDDT > 90, blue), (2) confident (70 < pLDDT ≤ 90, green), (3) low confidence (50 < pLDDT ≤ 70, yellow), and (4) very low confidence (pLDDT ≤ 50, red).

#### Table of Contents for SOM

#### Figure S1: Detail Page of TMVisDB Web Interface

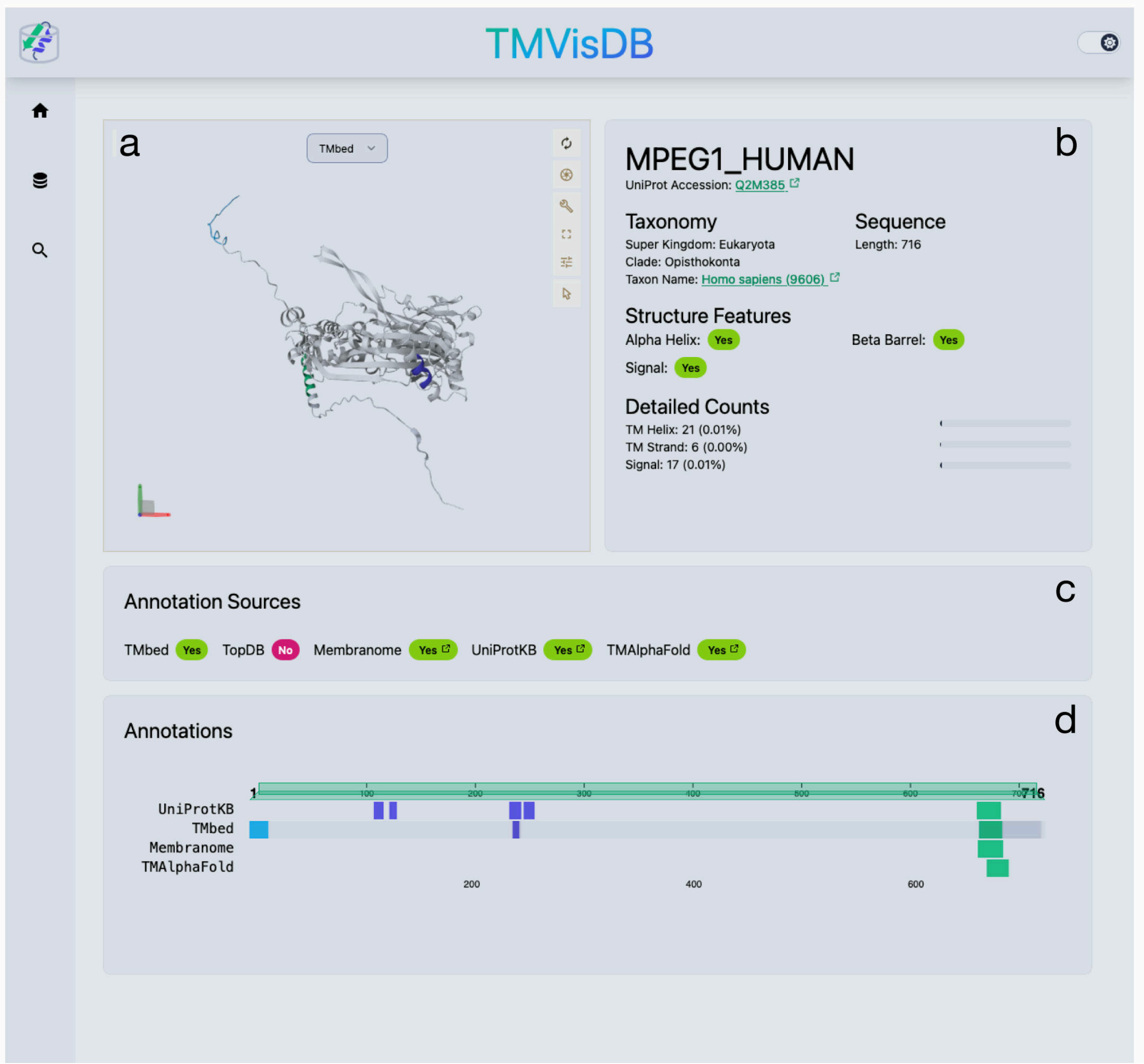

**Fig. S1**: **Detail Page of TMVisDB Web Interface for Q2M385.** Panel **a** presents the interactive AlphaFold2 [1] 3D structure viewer with customizable coloring schemes based on different annotation sources. Users can access detailed residue information, including type, structural annotation, and pLDDT score through atom-level hover interactions.

Panel **b** provides essential protein information from TMVisDB, including UniProtKB references, taxonomic data with relevant links, and a summary of TMbed-predicted structural features (signal peptides, transmembrane helices, and beta-barrels) [2].

Panel **c** compiles available protein annotations with direct links to their respective source databases.

Panel **d** features an interactive annotation viewer, built on EBI's nightingale components [3], enabling seamless integration with the 3D structure visualization through region highlighting and sequence navigation capabilities.

#### Figure S2: 3D structure and membrane topology visualization of Deoxyribonuclease I fragment

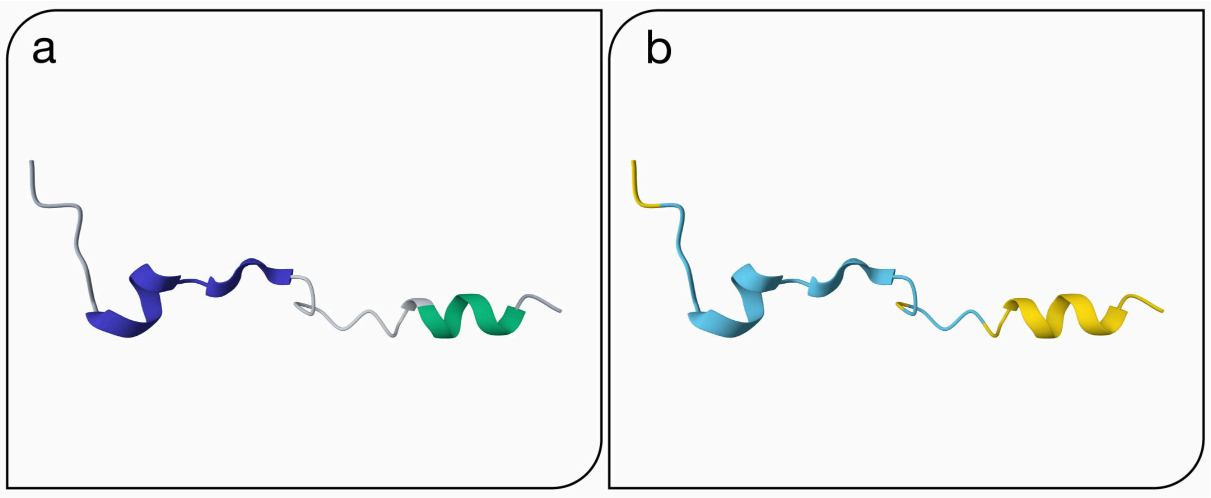

**Fig. S2**: **Deoxyribonuclease (DNase) I fragment (A0A192ZHB2)** Panel **a** displays the predicted transmembrane topology and Panel **b** shows the AlphaFold2 confidence scores as described above.

While TMbed predicts two transmembrane segments (one helix and one beta-strand) that align with structural elements in the AlphaFold2 model, their membrane-spanning arrangement appears structurally unlikely. The generally low pLDDT scores suggest limited confidence in the structural prediction, warranting careful interpretation of both TMbed and AlphaFold2 predictions for this protein.

#### Figure S3: 3D structure and membrane topology visualization of the Macrophage-expressed gene 1 protein

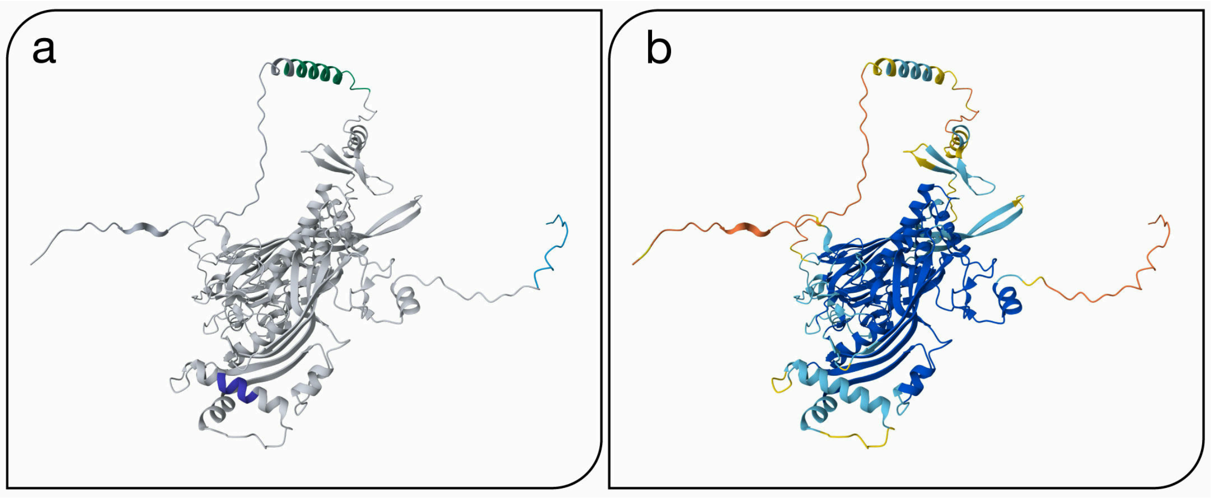

**Fig. S3**: **Macrophage-expressed gene 1 protein (Q2M385)** Panel **a** shows the predicted transmembrane topology and Panel **b** displays AlphaFold2 confidence scores as described above.

The predicted transmembrane helix (TMH) corresponds to a structural region with low to confident pLDDT scores. Notably, while TMbed predicts a single transmembrane beta-strand (TMB), AlphaFold2 suggests a confident helical structure in this region, indicating the TMB prediction is likely incorrect.

#### Figure S4: 3D structure and membrane topology visualization of DnaJ homolog subfamily C member 11

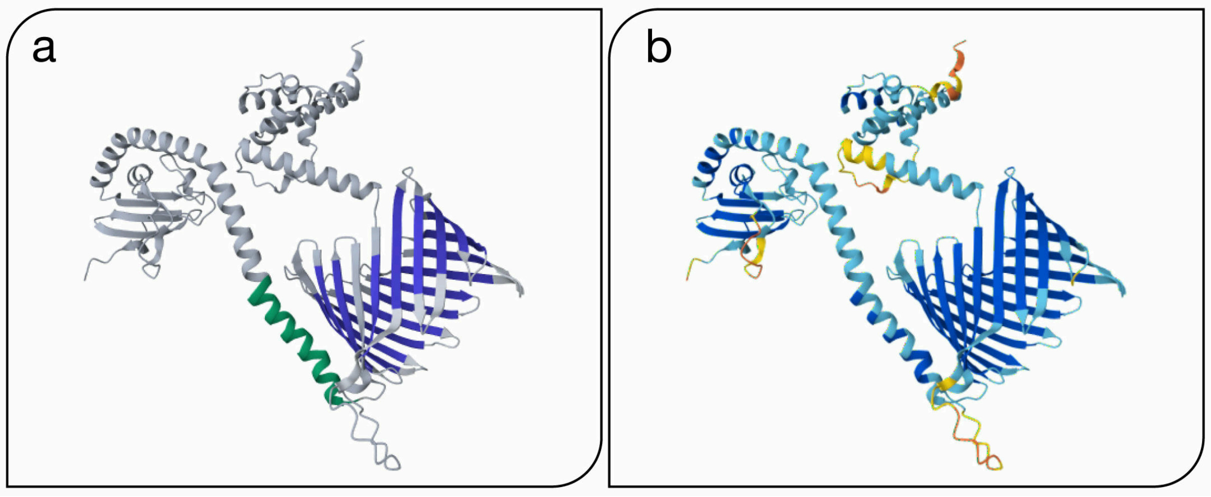

**Fig. S4**: **DnaJ homolog subfamily C member 11 (Q9NVH1):** Panel **a** presents the predicted transmembrane topology and Panel **b** shows AlphaFold2 confidence scores as described above.

The transmembrane predictions align well with the structural elements in regions of high pLDDT scores. Both the predicted alpha-helix and beta-barrel exhibit dimensions compatible with typical membrane thickness.

#### References - Supporting Material

[1] J. Jumper *et al.*, “Highly accurate protein structure prediction with AlphaFold,” *Nature*, vol. 596, pp. 583–589, 2021, doi: 10.1038/s41586-021-03819-2.

[2] M. Bernhofer and B. Rost, “TMbed: transmembrane proteins predicted through language model embeddings,” *BMC Bioinformatics*, vol. 23, p. 326, 2022, doi: 10.1186/s12859-022-04873-x.

[3] G. A. Salazar *et al.*, “Nightingale: web components for protein feature visualization,” *Bioinformatics Advances*, vol. 3, no. 1, p. vbad64, 2023, doi: 10.1093/bioadv/vbad064.
